# The interplay between cortical tracking and functional connectivity in musicians and non-musicians listening to speech and music

**DOI:** 10.64898/2026.10.27.762164

**Authors:** Noemi Bonfiglio, Christoforos Souganidis, Anastasia Klimovich-Gray, Nicola Molinaro

**Author notes:** Corresponding authors: Noemi Bonfiglio, Nicola Molinaro, Address: Mikeletegi Pasealekua, 69, 20009 Donostia-San Sebastiàn, Gipuzkoa, Spain.

## Abstract

Musical expertise is associated with auditory processing advantages which can extend to the speech domain. However, the origins of such domain-general benefits remain unclear. Recent frameworks propose that better rhythm processing in musicians might be at the basis of the observed advantages in processing music and speech. Auditory cortical tracking and inter-regional interactions between auditory and motor areas have been proposed as key neural mechanisms. In this study, we asked whether expertise-related increase in cortical tracking would be elicited by naturalistic stimuli in clear listening conditions, and whether such effects would be related to audiomotor functional connectivity and individual rhythmic abilities. We compared MEG data from expert musicians and controls with no musical playing experience while they listened to two-minutes long stories and excerpts of piano music. We also collected behavioral data in a rhythmic task testing speech (syllable) production. We found that musicians showed enhanced tracking and enhanced functional connectivity in both speech and music conditions, as well as better rhythmic abilities relative to non-musicians. Alpha connectivity between the right auditory cortex and inferior frontal gyrus negatively predicted cortical tracking across all participants in the speech condition, while no relation was observed for music. We suggest that stronger cortical tracking in musicians is due to domain-general differences in auditory sensory processes and is not driven by motor influences in our experiment. Moreover, a trade-off between cortical tracking and inter-regional communication might be at the basis of optimal perception of speech in both musicians and non-musicians in clear listening conditions.

## Introduction

Learning to play a musical instrument with high proficiency entails important neuroplastic changes at the anatomical and functional level [1,2], which have been associated with benefits in cognitive functions [3,4]. In the domain of auditory perception, studies have found more precise and efficient perceptual processing in musicians relative to non-musicians [5,6]. These benefits seem to extend to non-musical stimuli, including syllables and words [7–10] and continuous speech in noisy listening conditions [11,12]. Even if the robustness of this advantage is still under debate [13,14], there is now increasing interest in understanding the underlying cause of these effects. On the one hand, musicians might exhibit perceptual advantages due to better processing of the acoustic features of speech [6,7,15,16]. On the other hand, musical practice might affect perception in a more indirect way, by improving domain-general cognitive abilities [4,17,18]. Overall, to better understand the nature of musicians’ advantages, where present, and differences in processing more in general, it is crucial to complement behavioral studies with investigation of the underlying neural mechanisms.

One of the key proposed mechanisms of music-to-speech transfer effects is shared reliance on rhythmic processing [19,20]. In line with this view, a recent study found that the correlation between musical training (self-reported through the Goldsmiths Musical Sophistication Index questionnaire) and phonological language processing was mediated by rhythmic abilities, and in particular by the performance in the Spontaneous Speech Synchronization task (hereafter, “SSS”) [21], which measures the synchronization between participants’ speech production and a concurrently perceived syllable stream [22]. This suggests that both phonological perception and musical practice rely on developing robust rhythmic abilities, which can then facilitate perception of both speech and musical stimuli. At the neural level, high synchronization in the SSS was associated with stronger brain-stimulus phase-locking during syllable listening and with enhanced structural connectivity between auditory and frontal regions [22]. These findings align well with the Processing Rhythm In Speech and Music framework (PRISM) [20], proposing that musical practice can lead to transfer effects on speech processing through sharpening of three types of neural mechanisms: basic auditory processes, entrainment of neural oscillations to the rhythmic stimuli (i.e., speech and music), and interactions between sensory and motor neural processes.

Rhythmic or quasi-rhythmic stimuli generate a time-locking of neural activity that is considered especially relevant for speech processing [23]. However, the neural origins of such phenomenon are under debate, with some supporting a crucial role for the entrainment of endogenous oscillations [20,24,25] and others viewing it as a byproduct of evoked response to salient acoustic events [26,27]. Here, we will use the term “cortical tracking” to define time-locking in a way that is agnostic to this debate [23]. Within-domain effects of musical experience on cortical tracking of music-like stimuli has been previously observed [28–31]. However, the extent to which these effects can generalize to the speech domain is still unclear, as only few studies examined this question and reported mixed findings [12,29,32]. In the study of speech tracking, methodological choices might have a role in determining whether modulations from individual musical experience will emerge, including the music profile of the participants and the manipulations of the stimuli. Here, we test a sample of highly expert musicians, who most likely exhibit the neurofunctional changes associated with several years (>10 in our sample) of formal training and practice, as well as possible neural predispositions that made them more prone to pursue musical training and reach such a high level of musical proficiency. Further, we examine a relatively naturalistic condition (i.e., listening to narratives and musical pieces) to explore group differences in the fundamental properties of speech analysis.

A second proposed mechanism for rhythm processing is the integration between sensory and motor processes during perception [19], even in the absence of overt movements [33,34]. It has been proposed that stronger interactions between auditory and motor regions in musicians may be the main force underlying the observed speech processing advantage [35]. This “premotor” hypothesis builds upon the literature showing enhanced audiomotor connectivity in musicians [36–39], with functional differences being observed even during speech-related tasks [35,40]. How these structural and functional changes interact with cortical tracking remains an open question. In the general population, studies have shown that speech tracking can be modulated by oscillations in (pre)motor regions [41,42], which can provide precise temporal predictions to better align cortical activity to the incoming stimulus [43–46]. Predictive timing might be a fundamental aspect of rhythm processing mediating music-to-speech transfer effects. Some indirect evidence for this comes from the literature on motor synchronization tasks, in which participants tap along with a perceived beat or with a stream of rhythmic tones. Predictive behavior in tapping as well as tapping consistency were found to be correlated with musical experience [47], and musicians with earlier age of onset of musical training showed both increased gray matter volume in premotor regions and better tapping performance [48]. However, no studies to our knowledge have tested the possible interaction between cortical tracking of speech and music and motor influences in musicians.

The primary goal of this study was to assess whether professional musicians show different cortical tracking and functional connectivity patterns when tested with relatively naturalistic speech and music. We used magnetoencephalography (MEG) and measured cortical tracking through coherence, a well-established method which relies on phase alignment with the stimulus [49]. We hypothesized that, if musical expertise is associated with fine-tuning of general auditory tracking mechanisms, we would observe its effects on the phase alignment between the MEG signal and the stimuli across speech and music domains, even in the absence of stimulus manipulations. To put the “premotor” hypothesis at test, we complemented data on cortical tracking with analyses of functional connectivity, quantified as the weighted Phase Lag Index (wPLI, [50]) between activity in auditory and motor regions, a measure which had been previously used to study audiomotor connectivity during speech production with MEG [51]. If enhancements in auditory tracking are influenced by audiomotor interactions, as proposed by the “premotor” hypothesis, we would expect functional connectivity to be enhanced in musicians, and to positively predict cortical tracking across participants. Furthermore, we used a simple measure of rhythmic abilities in the speech domain to explore individual variability in behavior that might better characterize the neural differences between musicians and non-musicians. Our results showed that musicians exhibit enhanced tracking and enhanced functional connectivity as a group for both speech and music, but do not support the hypothesis that differences in cortical tracking are driven by motor influences in clear listening conditions.

## Results

### Stronger cortical tracking in musicians

24 professional musicians and 24 non-musician controls listened to fairy tales and pieces of classical piano music, presented in separate blocks. To ensure participants paid attention, they were asked to answer a two-choice comprehension question at the end of each story and a question about the author after each musical piece. Coherence between the segmented MEG sensor-level time series and the Hilbert transformed stimulus envelope was computed within the 0.5-10 frequency range, separately for speech and music stimuli. Coherence values were *z*-scored using a permutation procedure, and all participants showed significant cortical tracking above the permutation baseline (see “Brain-stimulus coherence” for details). We found that musicians exhibited stronger cortical tracking than non-musicians in the delta band for both speech (delta: 0.5-2 Hz; main effect of group: *F*(1, 46) = 4.98, *p* = .030; ƞ² = .10) and music (delta: 0.5-3.5 Hz; main effect of group: *F*(1, 46) = 4.78, *p* = .033, ƞ² = .09; Fig 1A). Moreover, in line with previous literature [52, 53], we found a significant right-lateralization of cortical tracking of speech stimuli in both the delta (main effect of hemisphere: *F*(1, 46) = 5.60, *p* = .022, ƞ² = .11; Fig 1B) and theta frequency bands (theta: 4-7 Hz; main effect of hemisphere: *F*(1, 46) = 6.89, *p* = .011, ƞ² = .13; S1A Fig).

**Fig. 1.**
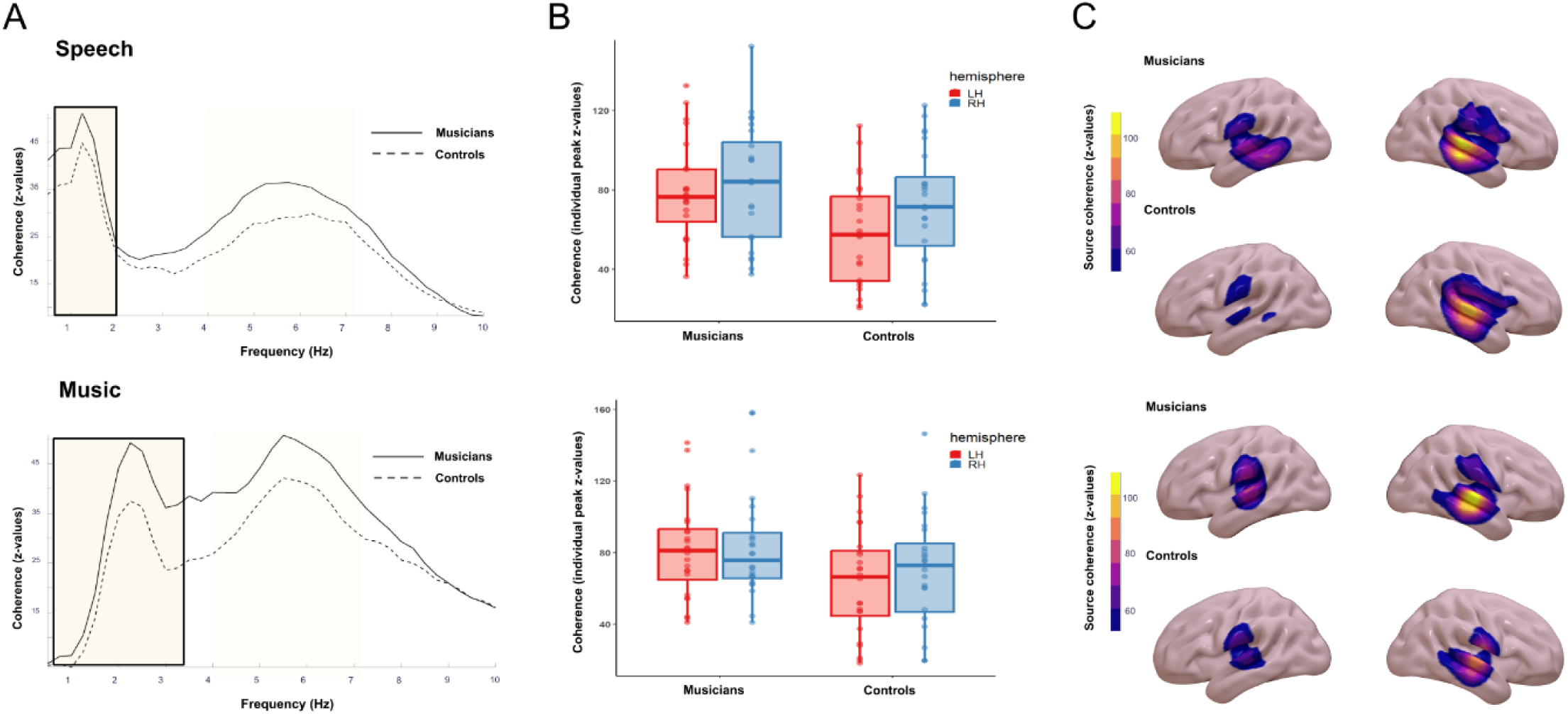
Brain-stimulus coherence in speech and music conditions. (A) Average *z*-scored coherence across participants in the 0.5-10 Hz range, averaged across 4 left and 4 right temporal virtual sensors. Shaded areas indicate the frequency ranges from which we extracted individual peaks for statistical analysis (delta in bold). (B) Boxplots of individual peaks of coherence in the delta band, on which we run the analysis. LH: Left Hemisphere; RH: Right Hemisphere. (C) Average source maps of *z*-scored coherence in the delta band. Boxplots and source maps for the theta band are shown in S1 Fig.

While differences in attention towards the stimuli have been associated with modulations in cortical tracking [54], behavioral responses do not support that the observed group differences are driven by attention, as musicians performed significantly better in music questions (*t*(44) = 3.08, *p =* .003, degrees of freedom reported after Welch adjustment; Musicians: mean = .67, sd = .18; Controls: mean = .49, sd = .22) but not in speech questions (*t*(44) = -0.14, *p =* .88; Musicians: mean = .77, sd = .13; Controls: mean = .78, sd = .11). Beyond *z*-scoring, we wanted to have an additional control that the effects really reflected cortical tracking and not power differences across groups. We ran the same statistical analyses on power values and we did not find any effect of group or hemisphere, confirming that our coherence effects are driven by phase synchronization with the stimuli (see “Control analyses of power” and S2 Fig for details). Finally, to confirm that our results were driven by activity in the auditory cortex, we computed coherence at the source level using the Dynamic Imaging of Coherence Sources (DICS) algorithm [55]. Since statistical analysis was already performed on the same data in sensor space, no further group-level statistical tests were performed on the source maps. We did, however, *z*-score the source-level coherence values to correct for coherence bias given by noise. This analysis revealed significant cortical tracking above the permutation baseline, which was localized, as expected, in bilateral Superior Temporal Gyrus, with a preference for the right hemisphere (Fig 1C). Table 1 shows the localization of the peaks of coherence at the group level in Montreal Neurological Institute (MNI) coordinates.

**Table 1.**
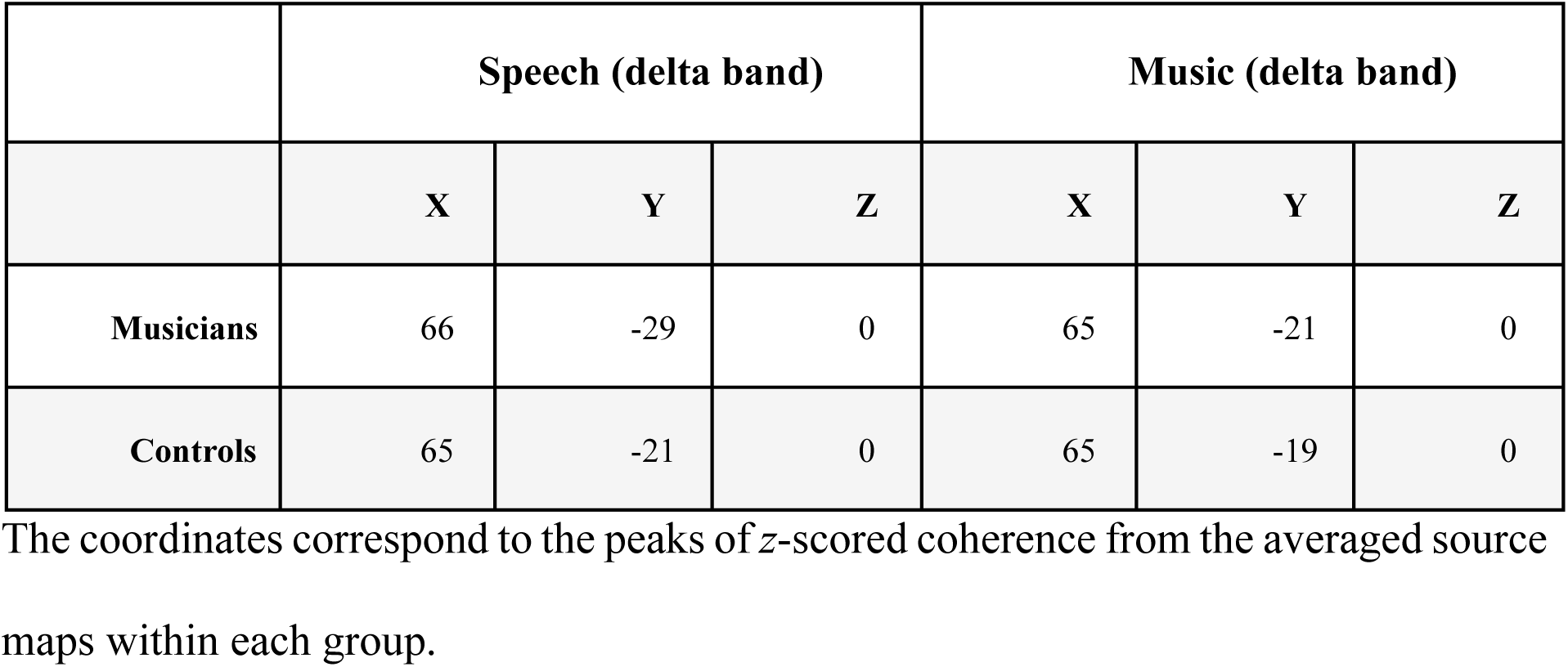
MNI coordinates of the peaks of coherence.

### Group differences in phase-based functional connectivity

To assess functional connectivity between auditory and motor regions during speech and music listening, we computed wPLI between source-level time courses in delta (0.5-2 Hz for speech; 0.5-3.5 Hz for music), theta (4-7 Hz), and alpha (8-12 Hz) frequency bands, based on the literature showing speech-related audiomotor connectivity in each of these bands [46,51,56]. We selected 2 auditory and 4 motor Regions of Interest (ROIs): Superior Temporal Gyrus, Transverse Temporal Gyrus, Precentral Gyrus, Middle Frontal Gyrus, Pars Triangularis and Pars Opercularis. One musician was excluded from subsequent analysis of connectivity as they consistently showed outlier values (> 3.3 standard deviation from the mean of the sample) in all frequency bands and conditions. Since our main goal was to explore group differences associated with musical expertise, we first identified the ROI connectivity pairs for which wPLI values differed between groups (|*t >* 2|; see “Statistical analyses”). Based on this criterion, we selected two ROI pairs for the speech condition (one in delta band, one in alpha band), and five ROI pairs for the music condition (four in theta band, one in alpha band), which all showed enhancements in musicians. Table 2 shows the seven selected ROI pairs.

**Table 2.** ROI connectivity pairs selected for statistical analyses.

| Stimulus condition | Hemisphere | Frequency band | Auditory ROI | Motor ROI | $t$ value<sup>a</sup> |
| --- | --- | --- | --- | --- | --- |
| Speech | Left | Delta | Superior Temporal Gyrus | Pars Opercularis | 2.30 |
|  | Right | Alpha | Transverse Temporal Gyrus | Pars Triangularis | 2.02 |
| Music | Left | Theta | Superior Temporal Gyrus | Precentral Gyrus | 2.73 |
|  | Right | Theta | Superior Temporal Gyrus | Middle Frontal Gyrus | 2.29 |
|  | Left | Theta | Superior Temporal Gyrus | Middle Frontal Gyrus | 2.23 |
|  | Left | Theta | Transverse Temporal Gyrus | Middle Frontal Gyrus | 2.34 |
|  | Right | Alpha | Transverse Temporal Gyrus | Middle Frontal Gyrus | 2.08 |

<sup>a</sup> $t$ values from two-samples Welch $t$ -tests comparing the groups. Positive $t$ values indicate musicians $>$ controls. No ROI pairs were found for which $t < -2$ .

Next, we evaluated whether the observed wPLI values were significantly different from chance by running surrogate tests for each participant, i.e., we compared them with surrogate distributions from signals with random phase relations but intact spectral properties. We then tested whether the results of the surrogate test differed between groups. We found that in the speech condition (Fig 2A), musicians were 10.06 times more likely to show significant delta connectivity between left Superior Temporal Gyrus - Pars Opercularis (odds ratio = 10.97, *CI* = 1.58-197.86, *p* = .039) and 3.6 times more likely to show significant alpha connectivity between right Transverse Temporal Gyrus - Pars Triangularis (odds ratio = 3.6, *CI* = 1.05-13.86, *p* = .049). This suggests that differences in significant phase-based connectivity, rather than in signal amplitude and Signal- to-Noise Ratio (SNR), were driving the difference identified in our initial ROI pairs selection (see “Statistical analyses” for details). In the music condition, we similarly found increased likelihood for musicians to show significant theta connectivity in two ROI pairs (Fig 2B), which however did not survive correction for multiple comparisons (left Superior Temporal Gyrus - Precentral Gyrus and right Superior Temporal Gyrus - Middle Frontal Gyrus; odds ratio = 3.97, *CI* = 1.20-14.48, *p* = .056 for both ROI pairs, fdr corrected for 4 comparisons).

**Fig 2.**
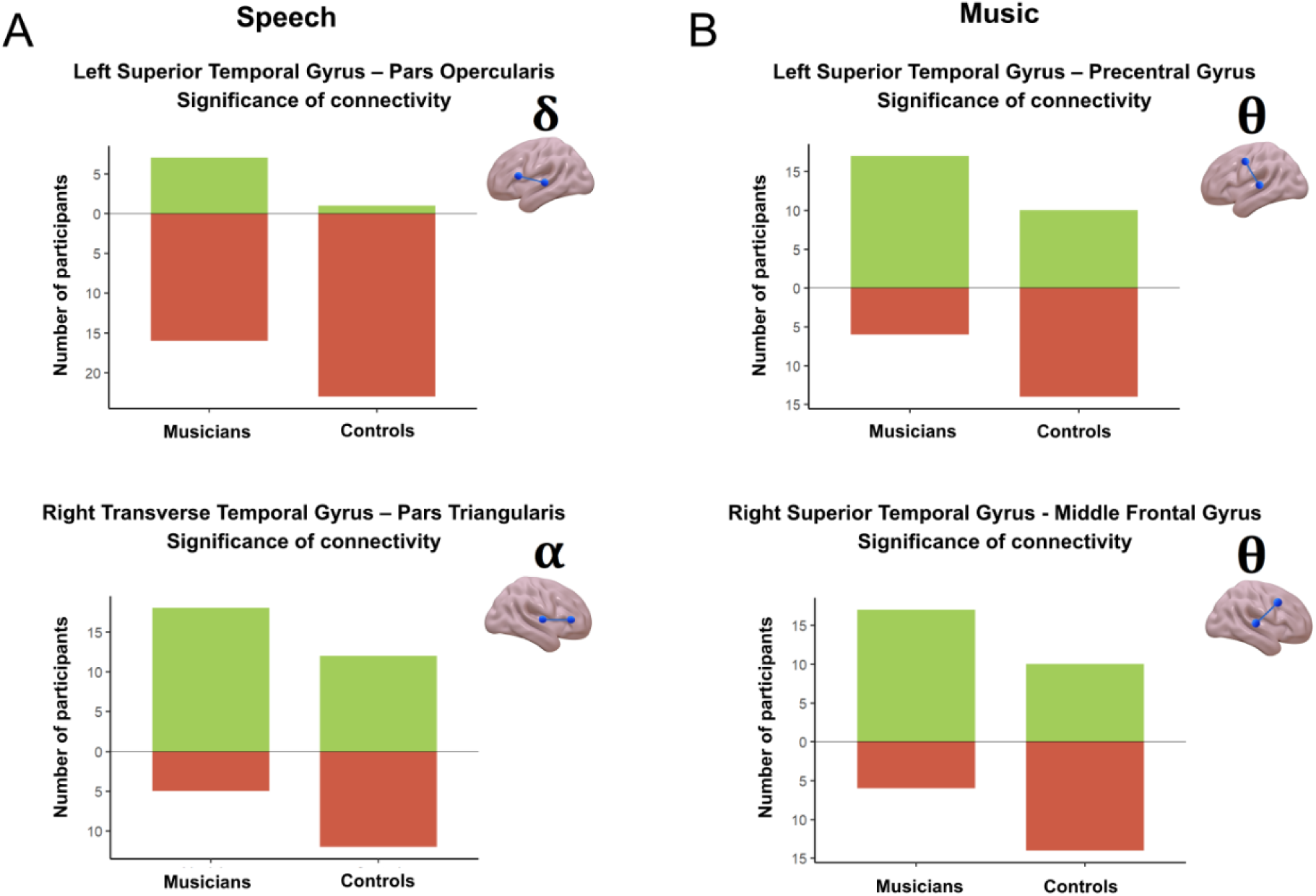
Group differences in phase-based connectivity. Barplots showing the number of participants with significant (green) and non-significant (brown) connectivity in the surrogate test, for speech (A) and music (B) conditions. Barplots for the remaining ROI pairs with no significant differences in the music conditions are shown in S3 Fig. ẟ: delta band; θ: theta band; *α*: alpha band.

### Better rhythmic abilities in musicians

To estimate participants’ rhythmic abilities in the speech domain we used a short steady repetition task. Participants were asked to listen to a 10-s stream of the syllable “ta” at a rate of 4.5 Hz and to reproduce it while matching the syllable rate as closely as possible. Analysis of the behavioral data was performed for 20 musicians and 18 controls due to data loss of the remaining audio recordings. We measured the performance in maintaining rhythm by estimating phase consistency of each individual production through a circular statistics approach [57]. In particular, we used the *z*-statistic from the Rayleigh test for circular uniformity as our measure of phase consistency. We considered consistency in relation to two reference rates: the rate cued by the stimulus (i.e., 4.5 Hz, Inter-Onset Interval, IOI, of 222 ms) and the average rate actually produced by participants. In this way, we estimated participants’ capacity to 1) extract and reproduce an external rhythm and 2) steadily keep any preferred pace, which could be different from the one cued by the task. We found that, on average, musicians outperformed non-musicians in both abilities (target rate: *t*(35) = 3.03, *p* = .004; individual rate: *t*(21) = 3.39, *p* = .002; degrees of freedom reported after Welch adjustment). As can be expected, across both groups maintaining any individual rhythm was easier and generated higher phase consistency values than reproducing the previously heard rhythm (Fig 3). We also observed different distributions of individual performance in maintaining the task rhythm and the individual rhythm, specifically in musicians (S4A Fig). For the task rhythm, musicians approximated a bimodal distribution with an almost-categorical distinction between good and bad performers. For the individual rhythm, on the contrary, all musicians showed a good performance, with less pronounced inter-individual variability. This indicates that even if some musicians were not able to faithfully reproduce the rhythm required by the task, they nonetheless consistently maintained a different, internally generated rhythm.

**Fig 3.**
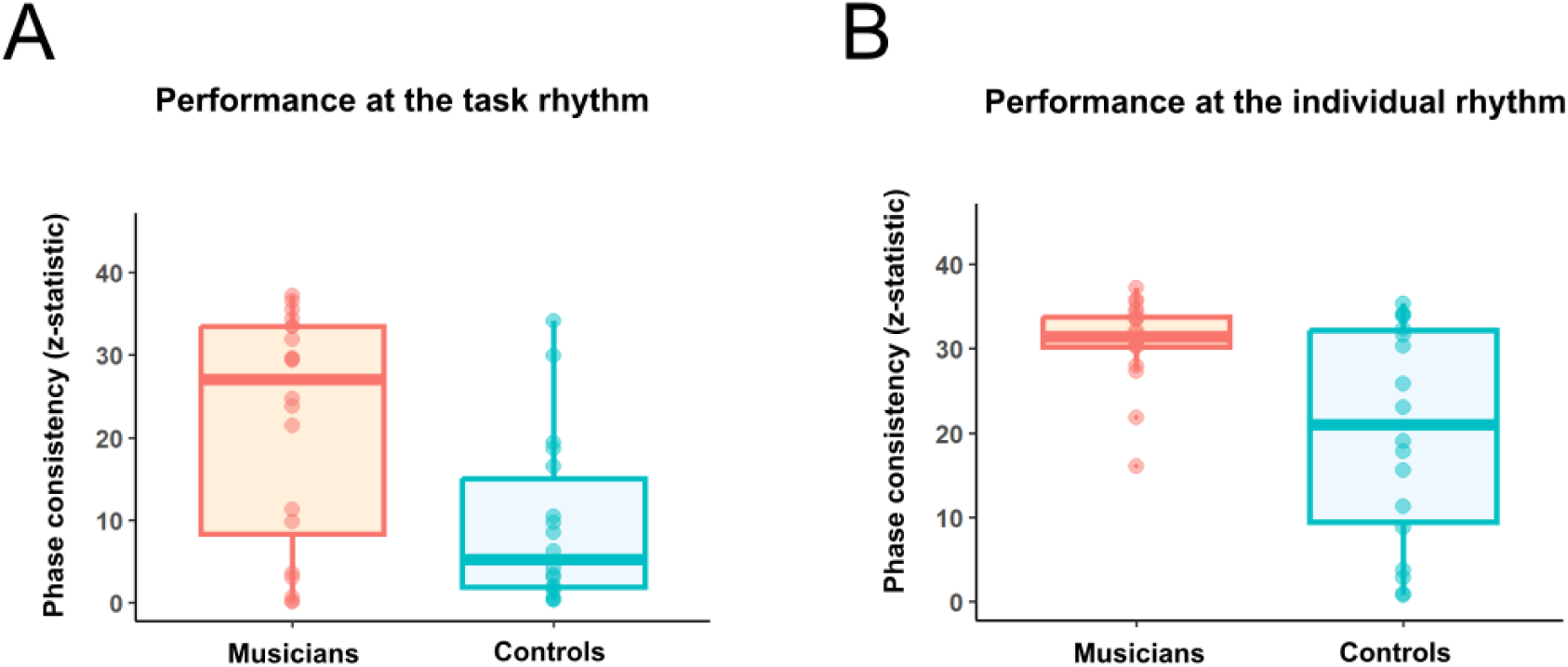
Individual performance in the steady repetition task. *Z*-statistic values from the Rayleigh test after computing phase relative to the task rate (4.5 Hz) (A) and the average individual rate (B) in the same trial. For both analysis and visualization, the trial showing the best performance in reproducing the task rate (out of 2 trials) was selected for each participant.

### Relation between cortical tracking, functional connectivity and rhythmic abilities

We hypothesized that, if tracking is facilitated by top-down modulations from motor regions in musicians, increased cortical tracking in the delta band (i.e., where we found the group difference) would be associated with increased functional connectivity across participants. To isolate phase-based connectivity, we removed the effect of amplitude and SNR from the wPLI values by *z*-scoring them with the corresponding surrogate distributions. Contrary to our expectations, in the speech condition we found a negative relation between tracking and connectivity: increased delta tracking in the right hemisphere was associated with decreased alpha connectivity between right Transverse Temporal Gyrus - Pars Triangularis (main effect of *z*-scored wPLI in predicting coherence: *t*(43) = -2.85, *β* = -12.82, *CI* = -21.91– -3.74, *p* = .007), and we did not find evidence that this effect differed between groups (interaction wPLI*group: *t*(43) =0.21, *β* = 0.93, *CI* = -8.16–10.01, *p* = .838; Fig 4A). In the music condition, we did not find significant relations between connectivity and cortical tracking. We observed, however, that musicians with stronger delta tracking in the left hemisphere tended to show stronger theta connectivity between left Transverse Temporal Gyrus - Middle Frontal Gyrus, while no such tendency was observed for controls (interaction wPLI*group: *t*(43) = -2.35, *β* = -7.71, *CI* = -14.33- -1.09, *p* = .094 fdr corrected for 4 comparisons; S3D Fig).

**Fig 4.**
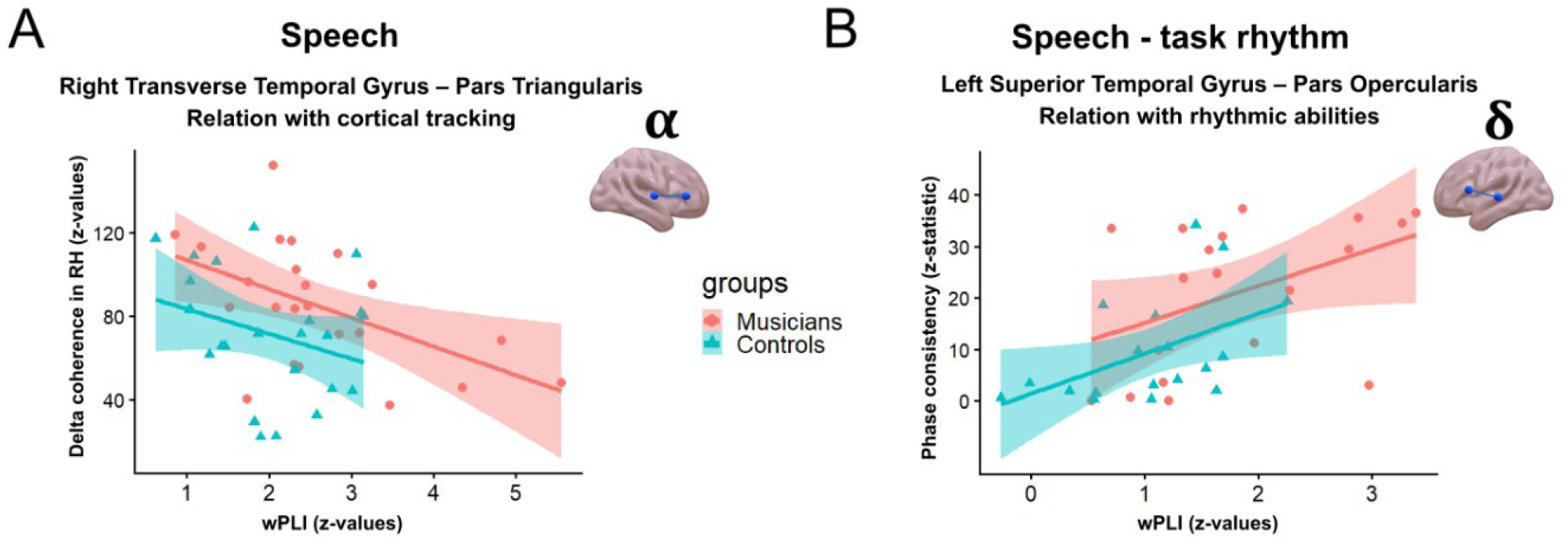
Relation between cortical tracking, functional connectivity and rhythmic abilities in the speech condition. (A) Relation between alpha connectivity (*z*-scored wPLI values) and cortical tracking in the delta band (*z*-scored coherence) in the right hemisphere. (B) Relation between delta connectivity (*z*-scored wPLI values) and performance in maintaining the task rhythm (phase consistency). Dots (musicians) and triangles (controls) represent individual participants.

Finally, we were interested in testing whether participants with better rhythmic abilities in the speech domain would show stronger cortical tracking of speech and functional connectivity during speech listening. While we did not find evidence for a relation with cortical tracking, we found that better performance in reproducing the rhythm cued by the task was associated with stronger delta connectivity between left Superior Temporal Gyrus - Pars Opercularis (main effect of *z*-scored wPLI in predicting phase consistency: *t*(33) = 2.83, *β* = 7.47, *CI* = 2.11-12.83, *p* = .031 fdr corrected for 4 comparisons), and we did not find evidence that this effect differed between groups (interaction wPLI*group: *t*(33) = 0.12, *β* = 0.33, *CI* = -5.03-5.69, *p* = .902; Fig 4B). Importantly however, we can not rule out that this finding is driven by differences in power and noise, as the majority of participants showed lower *z-*values than 2 (i.e., within two standard deviations from the mean of the surrogate distribution; in line with the results of the surrogate test, see Fig 2A).

## Discussion

In this study, we asked whether professional musicians show domain-general processing differences relative to non-musicians when listening to naturalistic speech and music stimuli. We evaluated two neural mechanisms which support rhythm processing across domains: cortical tracking and inter-regional communication between auditory and motor areas. Our results can be summarized as follows. First, while we found that musicians as a group show increases in the two processes across domains, we did not find evidence that cortical tracking is facilitated by communication with motor areas in our experiment, possibly because increased timing sensitivity in the musicians’ brain locates in the auditory cortex itself. Second, across all participants, cortical tracking of speech shows a trade-off with functional connectivity between auditory and speech motor areas, possibly reflecting individual differences in the way the brain reaches optimal speech perception. Third, rhythm production abilities are enhanced in musicians relative to non-musicians, with pronounced individual differences within the musicians’ group. We now turn to discuss each of these points in the following sections.

### Cortical tracking and motor engagement in musicians

We found that musicians exhibited domain-general enhancements in cortical tracking relative to non-musicians. This result diverges from previous research that failed to find an effect of music experience on cortical tracking of speech [29,32]. Our study differs from these previous investigations in fundamental aspects. First, our musician participants had an average of 25 years of experience and a professional level of musical proficiency. It may be that such extreme characteristics contributed to reveal effects of music expertise that are more difficult to observe in studies treating musical experience as a continuum [29]. Second, phase-based measures of cortical tracking might be more sensitive to these effects than methods based on response amplitude, as a previous study comparing musicians and non-musicians in amplitude and latency of neural response to speech envelope did not find differences [32]. Finally, our stimuli were not manipulated and were consistent in rhythm with a large corpus of speech and music recordings [58], while the effects of musical experience might be hindered when speech is more unnatural [29].

The observed group differences in cortical tracking may be interpreted within different theoretical frameworks. Within an oscillatory view [20,24,25], musicians’ brain may more flexibly entrain endogenous neural oscillations to external auditory inputs. Alternatively, under an evoked-based interpretation [26,27], neural activity in the auditory cortex may be more sensitive to fine temporal differences in the input, generating evoked responses which are more precisely aligned to the stimuli. Under either hypothesis, auditory tracking mechanisms might be aided by oscillations and/or activity in motor areas which encode precise temporal predictions [43–46]. However, our findings do not support the presence of such mechanisms in clear listening conditions, or at least not to the same extent for speech and music domains.

We found different functional connectivity patterns in the speech and music conditions. During speech listening, connectivity between the left auditory cortex and the speech motor cortex was significantly different from noise only in a minority of participants. By contrast, during music listening, significant connectivity between auditory cortices and motor areas was observed bilaterally in most participants (Figs 2 and S3). This divergent pattern reflects the difference between listening to speech and music: during speech comprehension motor simulations might not be needed if the input is clear and easy to understand, as opposed to difficult listening conditions [42,59–61]; on the contrary, listening to piano pieces likely involved the engagement of motor areas to covertly simulate the production of the melody [37,62,63], especially in musicians (see also S3D Fig). This might be reflected in a mixture of top-down motor influences and bottom-up activation from auditory to motor areas. In fact, an exploratory analysis using directed Phase Lag Index on a subset of connectivity pairs did not show preferences for either direction across participants (see “Exploratory analysis of directed connectivity” and S5 Fig).

Rather than being driven by motor influences, our cortical tracking results might be due to domain-general differences in auditory sensory processes, overlapping between speech and music domains [20]. In line with this hypothesis, it has been proposed that more consistent auditory brainstem response might underlie better phoneme discrimination in musicians [64,65]. More recently, a large-scale replication study failed to find an association between musical experience and sub-cortical encoding of speech sounds [14], leaving open the possibility that the effects observed in the literature are due to differences in primary and associative auditory cortices instead. In particular, it has been proposed that structural differences reflect pre-existing musical predisposition, while functional differences in auditory responses would be due to training [66]. While musical experience has been previously associated with neurofunctional changes [1,2], we can not exclude in our study that these neural patterns were already pre-existing in our participants prior to starting their training, and that they reached a professional level exactly because of their neural predisposition. In sum, while we can not claim that these effects are caused by musical training, we suggest that stronger auditory tracking in musicians reflects differences in basic auditory processes, as our data do not support that it is driven by audiomotor interactions in this experiment.

### Alpha-mediated inter-regional interactions in speech perception

The alpha band might have a special role in mediating speech-related inter-regional communication, and such communication channel seems to be modulated by musical experience. In our study, the majority of musicians (18 out of 23) showed above-chance alpha connectivity between right auditory cortex and Inferior Frontal Gyrus, against half of the non-musicians (12 out of 24; Fig 2A). This is in line with previous findings showing a positive association between self-reported years of musical training and alpha connectivity between the auditory cortex and a wide speech-related cortical network during naturalistic speech listening [40]. Further in line with our results, such relationship was particularly strong in the right hemisphere [40].

While musicians as a group show both enhanced tracking and enhanced connectivity relative to non-musicians, individual musicians who show stronger tracking tend to have weaker alpha connectivity in our study. This pattern could reflect a confounding effect of power: increased alpha power, associated with reduced attention to the stimulus [67], thus reduced tracking [54], may yield higher phase-connectivity estimates due to increased Signal-to-Noise-Ratio (SNR). However, to minimize this confound, we used a *z*-scoring procedure with surrogate distributions to isolate the effects of phase from SNR. To further confirm that our results reflect true phase-based connectivity in the alpha band, we performed an additional control analysis on alpha power at the frequencies used for connectivity analysis, and we did not find either differences between groups or correspondence between alpha power and functional connectivity estimates (see “Control analyses of power” and S6 Fig for details). Therefore, rather than being driven by attention and SNR, we suggest that our results reflect individual variability in how the brain processes speech in ideal listening situations.

Since our participants have normal hearing and optimal clear speech perception, cortical tracking and alpha mediated inter-regional interactions might provide complementary contributions to such perceptual efficiency. In this view, the observed trade-off between the two processes reflects normal variation in how perception is optimized given the individual’s neurobiological constraints and experience-dependent plasticity. Inter-regional interactions might be particularly beneficial for individuals whose less precise tracking of speech rhythms does not enable a rich sensory representation. By contrast, in individuals with more fine-tuned auditory tracking, these alpha-mediated interactions might be redundant and kept to a minimum, even generating non-significant phase-connectivity values. Because musical practice is associated with neural changes in both auditory sensitivity [5–8,66] and audiomotor interactions [35–40], as well as with differences in neural predisposition [66], musicians are more likely to position themselves at either extreme of this tracking-communication continuum. This helps explain the group differences we observe in both cortical tracking and alpha connectivity, and might potentially explain why previous investigations reported mixed findings on the relationship between musical experience and cortical tracking [12,29,32].

From our data, we cannot disentangle the precise functional role of alpha connectivity, but speculations can be made based on previous literature. For example, right-lateralized alpha connectivity during covert speech has been interpreted as a combination of top-down and bottom-up interactions that maintain a perceptual representation of the internally produced output [51]. In our study, this bidirectional communication might enhance the representation of perceived speech when it is not optimally encoded at the first stages of auditory analysis (see also “Exploratory analysis of directed connectivity” and S5 Fig). Moreover, greater engagement of right-lateralized inter-regional communication in musicians may support a more fine-grained processing of spectral cues [40], in line with the observed right-hemispheric bias for spectral processing during music perception [68]. These hypotheses are not mutually exclusive, and future studies using more controlled designs may explore them in greater detail.

### Individual differences in musicians’ rhythmic speech production

In our study, musicians showed better rhythmic abilities than non-musicians, but we did not observe a relation between behavioral data and cortical tracking. One explanation is that production data in this study was too limited, as we only considered 10 seconds of production from 38 participants. Moreover, speech listening likely involves several interacting aspects of rhythm processing, which might not be captured by the rhythmic task used in this study, relying on maintenance of the rhythm in working memory and production processes. Future studies using a more comprehensive characterization of rhythmic abilities might be able to better characterize the interplay between musical experience, rhythmic abilities and neural processing of speech.

Our phase-consistency method allowed us to explore different sub-processes in place in the steady repetition task. Performance around the individual rhythm showed little individual variability in musicians, suggesting an uniform and strong ability to keep a steady pace in production. Nevertheless, in performance around the task rhythm, about half of the musicians showed significantly lower values than the rest of the group, indicating a difficulty in maintaining and precisely reproducing the previously heard rhythm (Figs 3 and S4). The preferred rate of these bad performers musicians was still close to the target (S7A Fig), but even small deviations of less than 0.5 Hz generated low phase-consistency estimates. By contrast, among non-musicians, the distribution of preferred rate was more spread, indicating increased difficulty in approximating the rhythm cued by the task (S7B Fig).

Our finding of a bimodal distribution of rhythmic abilities in musicians resembles the results for the SSS task presented for the first time in Assaneo et al. [22]. These authors propose that the general population can be categorically divided in “high synchronizers” and “low synchronizers”, whereby some individuals spontaneously synchronize to the concurrently perceived speech stream, while others maintain their preferred rhythm without being influenced by it [22]. Our results suggest a more complex picture, as we show that a bimodal distribution can also be present in an explicit and delayed speech rhythm task, at least in a sample of professional musicians. In our study, the increased complexity of the steady repetition task relative to the SSS may have restricted the characteristic bimodal distribution to the group of participants who can precisely keep a rhythm (i.e., any preferred rhythm) throughout production. However, it remains an open question for future studies what determines the difference between good and bad performers among musicians.

### Limitations and outstanding questions

The use of naturalistic stimuli presented in clear listening conditions allowed us to examine the presence of domain-general effects when no additional compensatory processes or listening effort are involved. However, one fundamental limitation of this approach is that we can not draw strong conclusions on the relation between our neural results and auditory perception in terms of intelligibility and perceived clarity. Thus, while many studies in the literature support the presence of perceptual benefits in musicians [9–11,15, 16,18], our study can not provide evidence in this direction. Moreover, the relationship between cortical auditory tracking and speech intelligibility is not linear, and stronger tracking does not always correspond to better perception. For example, stronger speech tracking was observed in older adults and individuals with hearing impairment, possibly indicating a compensatory mechanism in these populations [69,70]. On the contrary, speech tracking was found to be lower in individuals with dyslexia [71,72] and positively related to individual phonological decoding in both dyslexic and non-dyslexic readers [72]. Thus, one outstanding question for future studies is whether the effects of musical expertise that we observe at the neural level underlie noticeable benefits in perceptual and phonological processing of speech.

Auditory speech tracking has been proposed to work as a temporal sampling mechanism to extract relevant information from the speech stream [73]. In this view, its disruption would be at the basis of the phonological deficits observed in individuals with dyslexia, and musical training might be a valuable tool to restore it [74]. In line with this, longitudinal studies have shown positive effects of musical training on reading and phonological abilities in dyslexic children and poor readers [75–77]. Our results on speech tracking in musicians add interesting evidence in this direction. However, our cross-sectional approach leaves open some fundamental questions. First, because all our participants are professionals with extensive musical experience, we can not tell what is the necessary amount of training to reach an increase in cortical tracking. Moreover, we can not disentangle between the effects of training and pre-existent neural differences reflecting predisposition. Our data further show that among musicians there is some variability in the strength of cortical tracking as well as in the engagement of inter-regional connectivity during speech listening. Therefore, an interesting question for future studies is what type of training and experience better predicts neural refinements in either of the two processes.

## Conclusions

While speech processing differences in musicians have been previously observed in difficult listening situations, we here show that they can be present also in easy listening conditions and with relatively naturalistic speech and music stimuli. Moreover, we extend previous work by showing a trade-off between cortical tracking and inter-regional communication during speech perception. Individual variability in the relative engagement of the two processes might explain why previous studies did not find consistent effects in musicians. Therefore, combining cortical tracking and functional connectivity methods may be a promising approach to better characterize these individual differences in future research.

## Materials and methods

### Participants

26 musicians who received formal musical training were recruited through collaboration with music schools and orchestras located in Donostia-San Sebastian. All of them had experience in playing at least one musical instrument for 10 years or more (mean = 29.4; sd = 14.6), reached a professional level of proficiency in at least one instrument and spent at least two hours per week playing, as reported through the Edinburgh Lifetime Musical Experience Questionnaire (ELMEQ; [78]). Notably, most musicians started formal musical training during childhood, and individual differences in years of experience are mainly due to differences in age between participants (see S1A Table for details). 2 musicians were excluded from the analysis due to noisy MEG recordings. The remaining 24 musicians were matched with the same number of control non-musicians by age (Musicians: mean = 43.1, sd = 16.7; Controls: mean = 43.3, sd = 16.9) and gender (females = 13 in both groups). Controls were recruited through the PARTICIPA database of the Basque Center on Cognition, Brain and Language (BCBL), and reported to have had very little musical training (mean = 1.6 years, sd = 2.4 years) and to not play any instrument at the time of the experiment. Musicians scored higher than non-musicians in the general score of the Goldsmiths Musical Sophistication Index questionnaire (Gold-MSI, [79]; Musicians: mean = 80, sd = 7.4; Controls: mean = 54.5, sd = 21.1) as well as in its subscales (S1B Table). All participants were right-handed native Spanish speakers, had no history of neurological illness and presented normal peripheral hearing. The BCBL ethical committee approved the experiment (REF: 040222SM; following the principles of the Declaration of Helsinki), and all participants signed an informed consent form.

### Stimuli and procedure

#### MEG experiment

Eight fairytale stories in Spanish and eight excerpts of classical piano music were used as speech and music stimuli for the MEG experiment (see S2 Table for the complete list). The stories were reduced and adapted from an open website (https://www.bosquedefantastias.com) and were recorded by a female Basque-Spanish native speaker who was instructed to read them with a natural prosody. The piano pieces were converted in audio format, reduced and adapted from the YouTube online platform. All stimuli were about 2 minutes long. The average modulation spectrum of the stimuli is consistent with the results by Ding et al. [58] on a large corpus of naturalistic speech and music recordings, showing peaks between 4-8 Hz and 0.5-3 Hz for speech and music respectively (S8 Fig).

The experiment was run using the Psychtoolbox [80] of Matlab (r2021b) [81]. Music and speech stimuli were delivered through plastic ear tubes in separate blocks, separated by a break, and in a randomized order. Order of the blocks was also randomized across participants. The whole experiment lasted about 40 minutes. Participants were seated in front of a black screen in which instructions appeared in white font at the beginning of each block, and a white fixation cross was displayed during stimuli presentation. For speech, participants were asked to listen to the story and answer a two-choice comprehension question at the end through bottom pressing. For music, they were asked to answer who was the author of the piece between two options.

#### Behavioral experiment

The behavioral experiment was performed in a separate session relative to the MEG experiment. Participants were seated in a sound-proofed cabin and the experiment consisted in a series of short tasks administered with the OpenSesame software [82]. Participants listened to auditory stimuli through headphones and their vocal responses were recorded through a microphone independent from the computer. For the purpose of this study, we analyzed the two repetitions of a steady repetition task, which were interleaved with other tasks that are not analyzed here. The two trials were not performed subsequently, as they belonged to different blocks of the experiment.

The task stimulus was a 10-second isochronous stream of the “ta” syllable with a duration of 222 ms (rate of 4.5 Hz). To create the stream, first a single repetition of the syllable was recorded by two native Basque-Spanish speakers, one man and one woman, who were also musicians. The recordings were then processed using Praat [83]: recordings were denoised, and the pitch was flattened and adjusted to 110 Hz for the male speaker and 220 Hz for the female speaker. Finally, syllable duration was adjusted to 222 ms, and 45 syllable repetitions were concatenated to reach the total duration of 10 seconds. During the experiment, instructions appeared in white font on the screen over a black background. Participants were asked to listen to the stream (matching their gender) and to reproduce it at the same pace. Right after the auditory presentation, the cue “repeat ta” appeared and stayed at the center of the screen for 10 seconds.

#### Structural MRI acquisition

In order to later perform source reconstruction of the MEG signal, participants’ high-resolution 3D structural MRIs (T1-weighted MP2RAGE sequence) were acquired with a 3 T Trio scanner (Siemens, Munich, Germany) and a 32-channel head coil. To limit head movement, the area between participants’ heads and the head coil was padded with foam and participants were asked to remain as still as possible. Snugly fitting headphones were used to dampen background scanner noise and to enable communication with experimenters while in the scanner. A total of 240 sagittal 1-mm thick contiguous images were acquired with the following parameters: repetition time (TR) = 5000 ms, echo time (TE) = 2.98 ms, inversion time (TI) = 2500 ms, flip angle = 5°, matrix size = 240 × 256, field of view (FOV) = 240 × 256 mm², and voxel size = 1 × 1 × 1 mm³.

### MEG data acquisition and preprocessing

MEG data were acquired in a magnetically shielded room using the whole-scalp MEG system installed at the BCBL. Due to the change of the system during the period of data collection, data from all the 24 musicians and 19 controls were acquired through an Elekta-Neuromag MEG system, and the remaining 5 controls with a Megin Triux Neo MEG system. Both systems were equipped with 102 sensor triplets (each comprising a magnetometer and two orthogonal planar gradiometers) that were uniformly distributed around the head of the participant. The head position inside the helmet was continuously monitored using five head-position indicator coils. The location of each coil relative to the anatomical fiducials (nasion, left and right preauricular points) was defined with a 3D digitizer (Fastrak Polhemus). Digitalization of the fiducials together with ∼300 additional points evenly distributed over the scalp of the participant were used to perform head movements compensation in the MEG data during preprocessing and to spatially align the MEG sensor coordinates with the T1 MRI brain images during the source reconstruction analysis. MEG recordings were acquired continuously with a bandpass filter at 0.01–330 Hz and at a sampling rate of 1 kHz. Heartbeat and eye movements were monitored with three pairs of electrodes each put in a bipolar montage: on the right chest and on the left back for electrocardiography (ECG); on the external chanti of each eye for the horizontal electrooculography (hEOG); above and below the left eye for the vertical (vEOG).

Spatiotemporal Signal-Space Separation (tSSS) [84], an extension of the signal space separation method [85], was applied offline to MEG data to subtract the environmental magnetic noise and correct for head movements, using the Elekta-Maxfilter software (r2.2) for the data acquired with the Elekta MEG system, and the mne.preprocessing.maxwell_filter function of the mne-python toolbox (r1.7.1) [86] for the data acquired with the Megin MEG system. Bad sensors (i.e., sensors presenting a flat or very noisy signal for most of the recording) were visually detected and interpolated during tSSS. Subsequent preprocessing was performed using mne-python. In particular, a band-pass FIR filter between 0.01 and 120 Hz was applied and ECG and EOG artifacts were detected and removed using independent component analysis. Finally, segments of signal that were noisy across most channels or that corresponded to periods of time in which the audio was slowed down due to a software error were removed. In particular, for 14 participants (9 musicians) an average of 3.76 % (range 0.5-6.7 %) of the data was removed.

### Brain-stimulus coherence

Coherence between MEG signal and stimuli envelopes was computed separately for speech and music conditions using the Fieldtrip toolbox (r2024-01-29) [87] in Matlab (r2021b). Stimuli envelopes were computed by applying the Hilbert transform to the broadband auditory signals. Once aligned, both audio and MEG signals were downsampled at 100 Hz and segmented into 4-seconds epochs, with 0.5 s of overlap between subsequent epochs. We first used the ft_freqanalysis function of Fieldtrip to compute the complex Fourier spectra of the signal between 0.5-10 Hz for the entire length of the data, using multiple tapers based on discrete prolate spheroidal sequences (DPSS) and a frequency smoothing of 1 Hz. We then computed coherence between MEG and audio signals using the ft_connectivityanalysis function on the complex Fourier spectra. This resulted in a value of coherence for each MEG sensor and for each 0.25-Hz-wide frequency bin in the 0.5-10 Hz frequency range.

We summed the coherence values for each couple of gradiometers to obtain the coherence of a new combined gradiometer (hereafter, “virtual sensor”). In particular, for each couple, we weighted the values according to sine and cosine at 20 different orientations, and we finally selected the orientation showing the maximum summed coherence across the two gradiometers. This procedure resulted in a total of 102 virtual sensors. Finally, to evaluate if coherence values signified above-chance cortical tracking, we performed *z*-scoring using permutations for each participant. We randomly scrambled the correspondence between the MEG signal and the audio across all the 4-seconds epochs, we computed the coherence between the MEG signal and the new random audios and combined each couple of gradiometers as done with the real coherence. This procedure was executed 100 times per participant and per condition and the resulting coherence was averaged across all virtual sensors in each permutation. Finally, to obtain the *z*-scored coherence values which were used in subsequent analysis, we subtracted the mean of the permuted distribution from the real coherence value at each virtual sensor and divided the result by the permuted distribution’s standard deviation.

Source reconstruction was performed by using the individual Freesurfer brain reconstructions (Freesurfer r7.3.2; [88]) from the structural MRI images, except for 4 participants (2 musicians) who did not undergo the MRI session and for whom the MNI template was used. Freesurfer reconstructions (hereafter, “MRIs”) were manually aligned with the MEG sensor coordinates using the mne.gui.coregistration function of mne-python. The resulting transformation coordinates were then imported and applied to the MRIs in Fieldtrip for source-level coherence computation. We computed volume-based source spaces with the ft_prepare_sourcemodel function, using a template model with a spatial resolution of 1 cm, warped on the individual MRIs or MNI template. We applied the ft_freqanalysis function to the downsampled and segmented data to compute the cross-spectral density (CSD) matrix across MEG and audio signals between 0.5-10 Hz, using multiple tapers based on DPSS and a frequency smoothing of 0.5 Hz. We then applied the ft_sourceanalysis function to the CSD matrix to compute source-level coherence between MEG and audio signals, using a DICS beamformer spatial filter with a regularization parameter (lambda) of 5 %.

As the group-level data at the sensor-level showed visibly more extended delta band peak for the music condition then for the speech condition, we chose slightly different delta ranges for this and subsequent analyses, considering 0.5-2 Hz for speech and 0.5-3.5 Hz for music, and a theta range of 4-7 Hz for both (Fig 1A). Coherence computation at the source level was restricted to a single frequency of interest within these bands. For each subject, we chose the frequency of interest separately for each band and condition based on the peak of coherence at the sensor level among all virtual sensors. As the peaks of coherence were also selected for sensor-level analysis (see Statistical analyses), this ensured correspondence between our sensor-level data and source brain maps. This procedure led to a total of four all-brain coherence maps for each subject. We then *z*-scored each of these maps separately following the same procedure used at the sensor level, computing the average coherence across all sources in each permutation. Finally, we averaged each of the four maps across participants within each group for visualization, while statistical comparisons were performed on the sensor-level data (see Statistical analyses).

### Audiomotor functional connectivity

To compute wPLI [50] between auditory and motor regions in delta, theta, and alpha frequency bands we performed source reconstruction of the raw (not segmented) MEG data in mne-python. The raw data was downsampled to 50 Hz and band-pass FIR filtered in each of the three bands. For consistency with the coherence analysis, delta band was defined as the range 0.5-2 Hz for the speech condition and 0.5-3.5 Hz for the music condition, theta band as 4-7 Hz and alpha band as 8-12 Hz for both conditions. A free-orientation forward model was computed using cortical-based source spaces based on the individual MRIs or MNI template with a spacing between grid points of ∼0.5 cm (‘oct6’), a one-layer boundary element model (BEM) with a spacing of ∼0.3 cm (‘ico5’), and transformation coordinates derived from manual alignment between individual MRIs and MEG sensor coordinates. Noise covariance was computed using resting state data filtered in the frequency bands of interest. Source time courses were then reconstructed using a vector Linearly Constrained Minimum Variance (LCMV) beamformer spatial filter [89] and reduced to scalar estimates by taking the norm across orientations. This procedure was performed separately for each filtered version of the signal and for speech and music conditions.

Six ROIs in each hemisphere were defined according to Freesurfer’s parcelation based on the Desikan-Killiany atlas [90]: Superior Temporal Gyrus and Transverse Temporal Gyrus (hereafter, “auditory ROIs”), Precentral Gyrus, Middle Frontal Gyrus, Pars Triangularis and Pars Opercularis (hereafter, “motor ROIs”). We extracted a time course for each ROI using the “pca flip” method, which consists in extracting the first singular vector through singular value decomposition, and then scaling and sign-flipping it. We then segmented the obtained signal into 4-seconds epochs and computed the average wPLI across epochs between the time course of each auditory ROI and each motor ROI in the same hemisphere with the mne-connectivity package (r0.7.0) [91] of mne-python. We chose to focus on within-hemisphere interactions as we hypothesized that they would be particularly relevant for cortical tracking in auditory cortices, but we do not exclude a potential role of inter-hemispheric communication, which could be explored in future studies. The wPLI computation resulted in a total of 16 ROI pairs across the two hemispheres, with corresponding wPLI values for each 0.25-Hz-wide frequency bin in delta, theta and alpha bands, separately for speech and music conditions.

### Rhythmic abilities

Analysis of the behavioral task was performed for 20 musicians and 18 controls due to data loss of the audio recordings for the remaining 10 participants. We first manually cut the segments of interest, i.e., the two repetitions of the steady repetition task, from the audio recordings collecting the whole behavioral experiment. The resulting audios were further segmented to remove occasional breaths, to ensure that results would not be biased by the breathing capacity of the participants (i.e., worse estimates of rhythmic abilities for participants who paused to breathe). Annotation of the onset times of each produced syllable was performed following a semi-automatic procedure. First, we converted to mono format and resampled the audio at 22050 Hz. Then, we used the Librosa python package [92] to extract an onset strength envelope based on the mel-based spectrogram and automatically detect the time points corresponding to peaks in the envelope, which roughly correspond to the occurrences of the vowel “a” in the syllable “ta”. For each peak time point, we established the preceding 50 ms as the time window to search for the syllable onset, which was mathematically defined as the first point in the window with amplitude greater than 100 times the mean amplitude in the preceding 10 ms. In this way, onsets were defined according to the relative change from a period of silence. The resulting time points were collected into TextGrid files for each audio segment, and were manually checked and corrected using Praat.

To estimate phase consistency, onset time points were converted to degree units in a circular scale, with the target beat aligned at 0°, considering the target beat time points that would be produced if participants were perfectly isochronous at the target rate. For example, for a target rate of 4.5 Hz and a corresponding Inter-Onset Interval (IOI) of 222 ms, the first five expected time points would be at 0, 222 ms, 444 ms, 666 ms, 888 ms. We converted each time point by subtracting the corresponding expected time point, dividing the result by the IOI and multiplying it by 360. We performed this procedure for the rate cued by the stimulus (i.e., IOI 222 ms, 4.5 Hz) and the one actually produced by participants (i.e., the mean IOI in the produced stream). The degree points were obtained separately for segments separated by breaths, and were then concatenated to obtain a single measure for the corresponding trial. Finally, we computed the *z*-statistic from the Rayleigh test for circular uniformity through the pycircstat2 python toolbox. For the task rhythm, we selected the higher *z* values across the two trials for each participant (i.e., considering the best performance across trials); for the individual rhythm, we selected the *z* values corresponding to the same trial. We then compared the groups in two-tailed two-samples Welch *t*-tests, separately for the target rate and the individual rate.

### Statistical analyses

#### Sensor-level coherence

All statistical analyses were performed in the R software (v4.5.0) [93]. For statistical comparison of sensor-level coherence, we selected the virtual sensor showing the highest *z*-scored coherence value for each participant in each hemisphere and across all frequency bins in delta and theta frequency bands (as defined in the Brain-stimulus coherence section). This was done to account for individual differences in peak sensor location, which heavily depends on head’s shape and position within the MEG helmet. As literature shows that tracking in delta and theta bands reflect different functional processes in the brain [53,94–96], and because our naturalistic speech and music stimuli were not acoustically matched (see Stimuli and procedure), we statistically tested each frequency band and each stimulus type separately. We run four two-way mixed-design ANOVAs (delta-speech; theta-speech; delta-music; theta-music) testing the effect of group (between-factor) and hemisphere (within-factor) on individual *z*-scored coherence peaks. Prior to testing, we checked the assumptions of ANOVA following Tabachnick and Fidell’s [97] indications. No statistical outliers (i.e., values that deviates from the mean of the whole sample by more than 3.3 standard deviations) were found in the data. We tested homogeneity of variance by using Levene’s test and found no violation of the assumption. Finally, we assumed normality of sampling distribution based on degrees of freedom for error, which was greater than 20 for each of our tests.

#### Functional connectivity

For each subject, we extracted the higher wPLI values across frequency bins for each of the 16 ROI pairs in each frequency band and condition. One musician was excluded from this analysis as they consistently showed outlier values (> 3.3 standard deviation from the mean of the sample) in all frequency bands and conditions. To explore differences associated with musical expertise, we restricted our statistical analyses on the connectivity ROI pairs showing group differences according to a normal approximation (|*t* > 2| in two-samples Welch *t*-tests; Table 2). We used surrogate tests to evaluate whether the observed wPLI values for each subject exceeded the significance threshold when compared with wPLI distributions from signals with random phase relations and intact spectral properties (for reference using the method for this purpose, see [51]). This strategy ensures that significance is driven by the presence of a phase relation between signals (i.e., true phase-based connectivity) rather than by spectral properties such as power and SNR. This is important to ensure that group-level results are driven by phase-based connectivity rather than by power differences between groups, as signals with higher SNR would allow for better phase estimation and generate higher wPLI values.

For each subject and for each of the selected connectivity pairs, we created 1000 surrogate signal pairs through the Amplitude Adjusted Fourier Transformed (IAAFT) algorithm [98]. This algorithm consists of an initial randomization which alters both phase relationships and power spectrum, followed by iterative spectral and amplitude adjustments until both Fourier coefficients and amplitudes match the original time series. We used the open-source python implementation of the IAAFT algorithm, which can be found at: https://github.com/mlcs/iaaft. For each subject and connectivity pair, we applied the IAAFT algorithm to the time course of each ROI independently. We then segmented the surrogate signals in 4 seconds epochs and computed wPLI values in the frequency range of interest as done for the real signals. We repeated this procedure 1000 times to generate a distribution of surrogate wPLI values for each 0.25-Hz-wide frequency bin.

For each subject and connectivity pair, the surrogate test evaluated whether the true maximum wPLI value fell beyond the 95th percentile of the corresponding surrogate distribution at the same frequency bin. We then tested whether musicians and controls had the same probability of showing significant phase-based connectivity by fitting logistic regression models with significance (yes/no) as dependent variable and group (controls/musicians) as predictor (*significance ∼ group*). We used treatment coding for the group variable and controls as the reference group to obtain odds ratios for musicians. As functional connectivity in distinct frequency bands are derived from differently filtered signals and constitute distinct communication channels [46,99–101], we considered each frequency band and condition separately, applying the fdr method for multiple comparison correction when testing multiple connectivity pairs in the same frequency band and condition.

#### Relation among neural and behavioral measures

To evaluate relations between our measures of functional connectivity, cortical tracking and rhythmic abilities, we ran multiple linear regression models. We considered the *z*-scored peaks of coherence in the delta band extracted in the previous analysis as our measure of cortical tracking. For functional connectivity, we derived a continuous measure of the strength of connectivity for each pre-selected ROI pair (Table 2) by *z-*scoring the real wPLI values, i.e., subtracting the mean and dividing by the standard deviation of the corresponding surrogate distribution. This procedure removed the possible confounding effect of spectral properties, which were common between the real and surrogate signals. Finally, for rhythmic abilities, we used the best *z-*statistic values across the two trials for each participant (i.e., considering their best performance) around the task and the individual rhythm. In each of the following analyses, we used sum contrast coding for the group predictor, with controls as the first level and musicians as the second level of the variable.

First, following our main hypothesis, we evaluated the relation between cortical and functional connectivity. Linear regression models had *z*-scored coherence in delta band as dependent variable, and *z*-scored wPLI (continuous), group (controls/musicians) and their interaction as predictors (*coherence ∼ wPLI*group*). Each of the seven connectivity ROI pairs was used to predict coherence in the same hemisphere and condition. As above, we corrected for multiple comparisons with the fdr method when testing multiple connectivity pairs in the same frequency band and condition.

Next, we fitted linear regression models to predict individual rhythmic abilities from delta coherence in the speech condition. As we had two behavioral measures (performance around the task rhythm and the individual rhythm) and two measures of coherence (one for each hemisphere), we ran a total of four regressions. We used the *z-*statistic values as dependent variable and used *z*-scored coherence (continuous), group (controls/musicians) and their interaction as predictors (*z-statistic ∼ coherence*group*). We did not report corrected *p*-values as none of these regressions were significant.

Finally, we asked whether functional connectivity during speech listening could predict rhythmic abilities. We ran regression models to predict the two behavioral measures from the two connectivity pairs in the speech condition (one left-lateralized in delta band; one right-lateralized in alpha band; see Table 2), for a total of four regressions. We used *z-*statistic values as dependent variable and used *z*-scored wPLI (continuous), group (controls/musicians) and their interaction as predictors (*z-statistic ∼ wPLI*group*). We used the fdr method to correct for multiple comparisons across the four regressions.

### Control analyses of power

We computed power spectra in the 0.5-20 Hz frequency range in Fieldtrip applying the Fast Fourier Transform method and a boxcar window taper to the same sensor-level segmented data used for the coherence analysis. This resulted in individual power maps with the same frequency resolution as coherence and functional connectivity, i.e., 0.25-Hz-wide frequency bins. Then, we linearly combined the power values for each couple of gradiometers as described for the coherence analysis to obtain power values for 102 virtual sensors.

First, to confirm that our cortical tracking effects reflected real phase synchronization with the stimuli and not power differences between groups, we extracted power values in delta and theta bands at the same virtual sensors and frequency bins used for coherence analysis, i.e., where each participant showed the highest value of coherence in each hemisphere in the delta (0.5-2 Hz for speech, 0.5-3.5 Hz for music) and theta (4-7 Hz) frequency bands. With this data, we ran the same ANOVAs which showed significant coherence effects, and found no effects of group (delta band for speech *F*(1, 46) = .39, *p* = .533; music *F*(1, 46) = .30, *p* = .581) or hemisphere (speech in delta band F(1, 46) = .186, *p* = .667; theta band *F*(1, 46) = .166, *p* = .684). Individual values for this analysis are shown in S2 Fig.

Then, to exclude that our alpha connectivity results were driven by power differences among participants, we examined alpha power in right temporal sensors. In particular, for each participant we extracted a power value in alpha band (8-12 Hz) selecting the right temporal virtual sensor showing the highest power in the frequency bin used for connectivity analysis, i.e., where each participant showed the highest wPLI value. First, we compared the two groups in alpha power and found no group differences (*t*(46) = -0.411, *p* = .682; degrees of freedom reported after Welch adjustment; S6A Fig). This contrasts with the pattern observed in alpha connectivity, where musicians were more likely to show supra-threshold values and, consistently, showed higher *z-*scored wPLI values as a group (*t*(39) = 2.268, *p* = .028; S6B Fig). Finally, upon visual inspection, we did not observe any relation between *z-*scored alpha connectivity and alpha power values, confirming that the *z-*scoring procedure effectively removed the effects of individual variability in signal power (S6C Fig).

### Exploratory analysis of directed connectivity

For a subset of connectivity ROI pairs, we conducted an exploratory analysis of directed connectivity using the directed Phase Lag Index (dPLI). This measure is conceptually similar to the wPLI, but instead of estimating the consistency of the phase lag between two signals, it estimates the direction of such lag, complementing the information provided by the wPLI. dPLI ranges from 0 to 1, with values between 0.5-1 indicating that the first signal is more likely to lead the second signal, and values between 0-0.5 indicating that it is more likely to lag. Because in our study the first “reference” signal is the one corresponding to auditory cortices, values between 0.5-1 reflect bottom-up interactions and values between 0-0.5 reflect top-down interactions.

As our main research question concerned the influence of functional connectivity on cortical tracking, we selected for this analysis the connectivity ROI pairs showing relations with coherence: Right Transverse Temporal Gyrus – Pars Triangularis in alpha band for speech; Left Transverse Temporal Gyrus – Middle Frontal Gyrus in theta band for music (which showed a non-significant tendency for the interaction between wPLI and group). We extracted source time courses from these ROIs with the “PCA-flip” method, we segmented into 4 seconds epochs as we did for the main analysis, and we computed the average dPLI values across epochs using the mne-connectivity package. Again, this resulted in one value for each 0.25-Hz-wide frequency bin in alpha (8-12) and theta (4-8) bands for speech and music respectively. Then, for each participant we selected the dPLI value corresponding to the frequency bin that was used in the main analysis, i.e., where each participant showed the highest wPLI value. Upon visual inspection, we observed that all participants showed dPLI values close to 0.5, ranging from 0.4 to 0.6 (S5 Fig). We thus concluded that our data show no preference for either bottom-up or top-down direction.

## Acknowledgements

We thank Laura Fernández-Merino, Jiaqi Mao and Maite Kaltzakorta for their help in data collection, Sara Azcarate for helping in correcting syllable onsets’ time stamps from the recordings of the rhythmic task, Mikel Lizarazu for advice on coherence computation and *z*-scoring, and Tomas Lenc for useful feedback on a previous version of the manuscript. We thank the Musikene music school for help in participants’ recruitment and the Donostia International Physic Center for providing the computational resources used to generate the surrogate data.

## Author contributions

**Conceptualization:** Noemi Bonfiglio, Christoforos Souganidis, Anastasia Klimovich-Gray, Nicola Molinaro.

**Data curation:** Noemi Bonfiglio, Christoforos Souganidis.

**Formal analysis:** Noemi Bonfiglio.

**Funding acquisition:** Nicola Molinaro.

**Investigation:** Noemi Bonfiglio.

**Methodology:** Noemi Bonfiglio, Christoforos Souganidis, Anastasia Klimovich-Gray, Nicola Molinaro.

**Project administration:** Nicola Molinaro.

**Supervision:** Nicola Molinaro.

**Visualization:** Noemi Bonfiglio.

**Writing - original draft:** Noemi Bonfiglio.

**Writing - review and editing:** Noemi Bonfiglio, Christoforos Souganidis, Anastasia Klimovich-Gray, Nicola Molinaro.

