## Supplementary figures and tables for "The interplay between cortical tracking and functional connectivity in musicians and non-musicians listening to speech and music"

### **Supplementary materials**

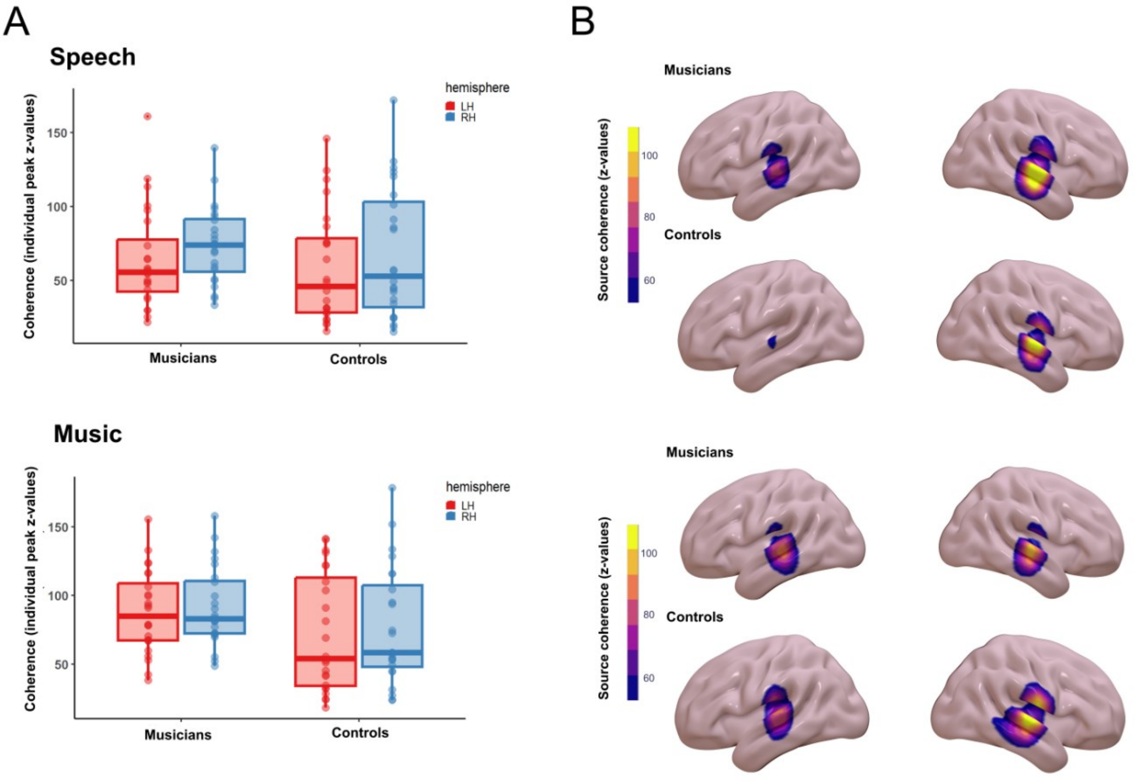


**S1 Fig. Brain-stimulus coherence in theta band.** (A) Boxplots of individual peaks of coherence. (B) Average source maps of *z*-scored coherence.

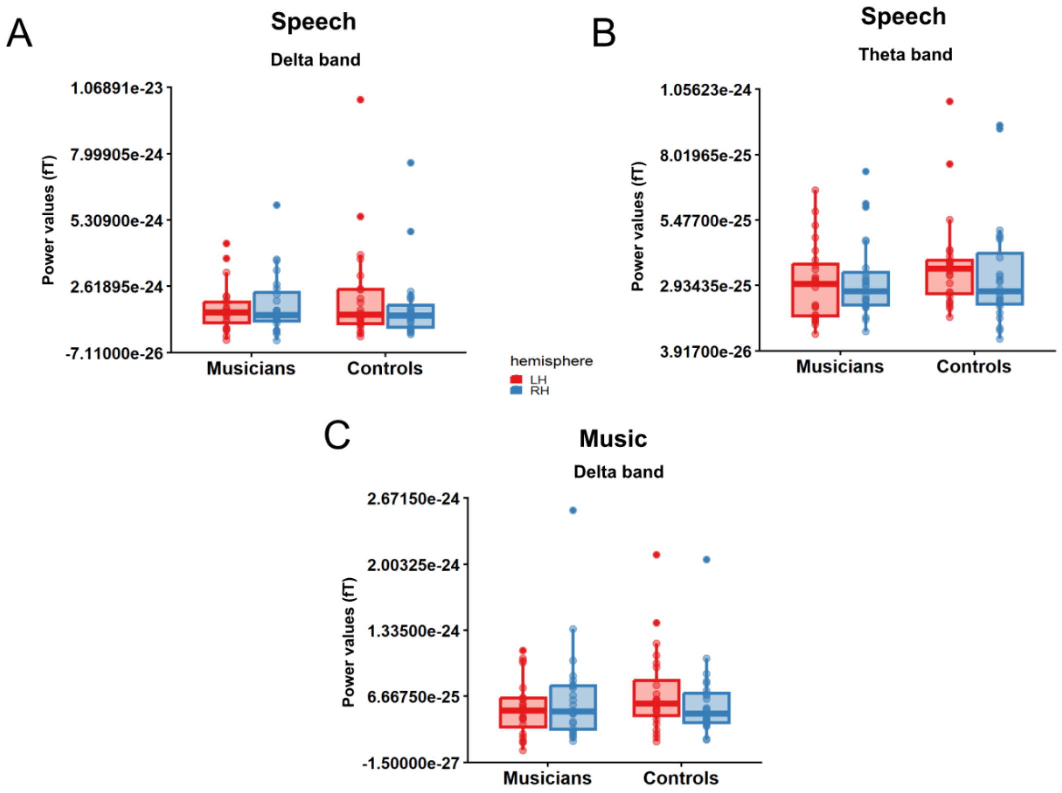


**S2 Fig. Power in delta and theta bands.** Values (here in femtotesla, fT) were extracted from the same virtual sensors and frequency bins used for coherence analysis, in the frequency bands showing significant effects: delta and theta for speech (A, B) and delta for music (C).

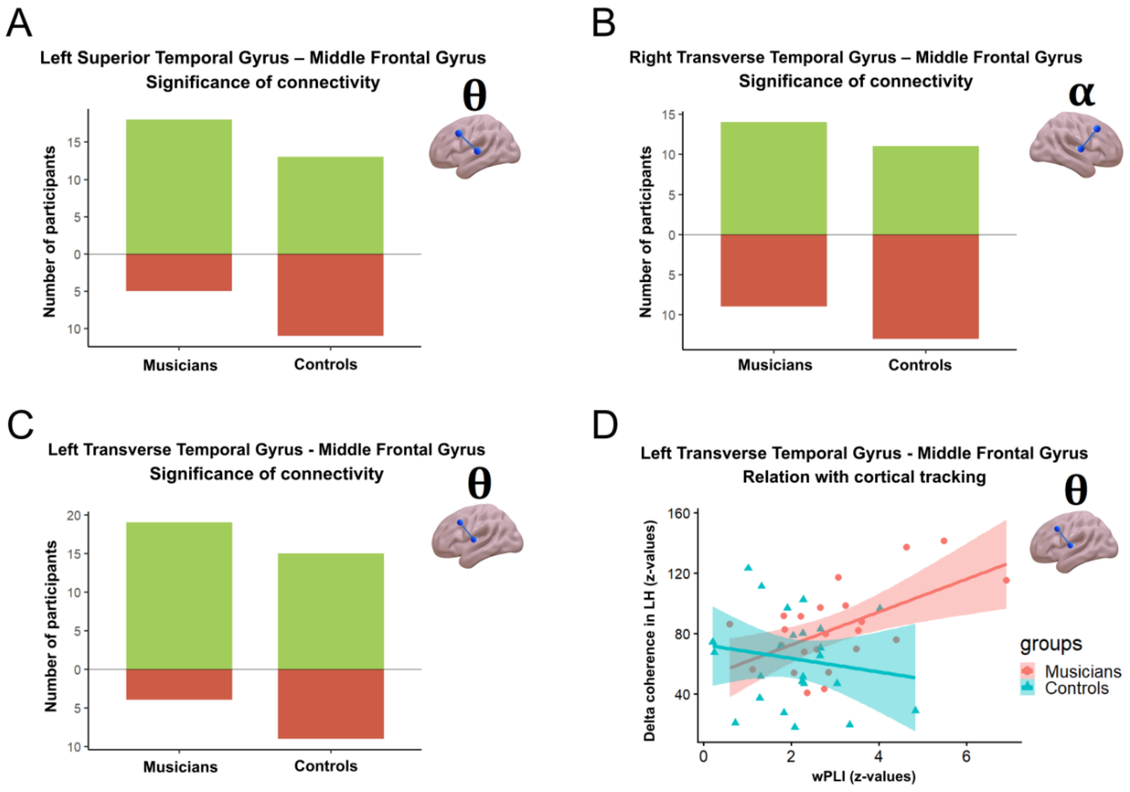


**S3 Fig. Non-significant effects in functional connectivity in music condition.** (A-C) Barplots showing the number of participants with significant (green) and non-significant (brown) connectivity in the surrogate test. Logistic regressions did not reveal significant differences between groups in these ROI pairs. (D) Relation between theta connectivity ($z$-scored wPLI values) and cortical tracking in delta band ($z$-scored coherence) in the left hemisphere (interaction wPLI*group in predicting coherence: $p$ = .094 fdr corrected for 4 comparisons).

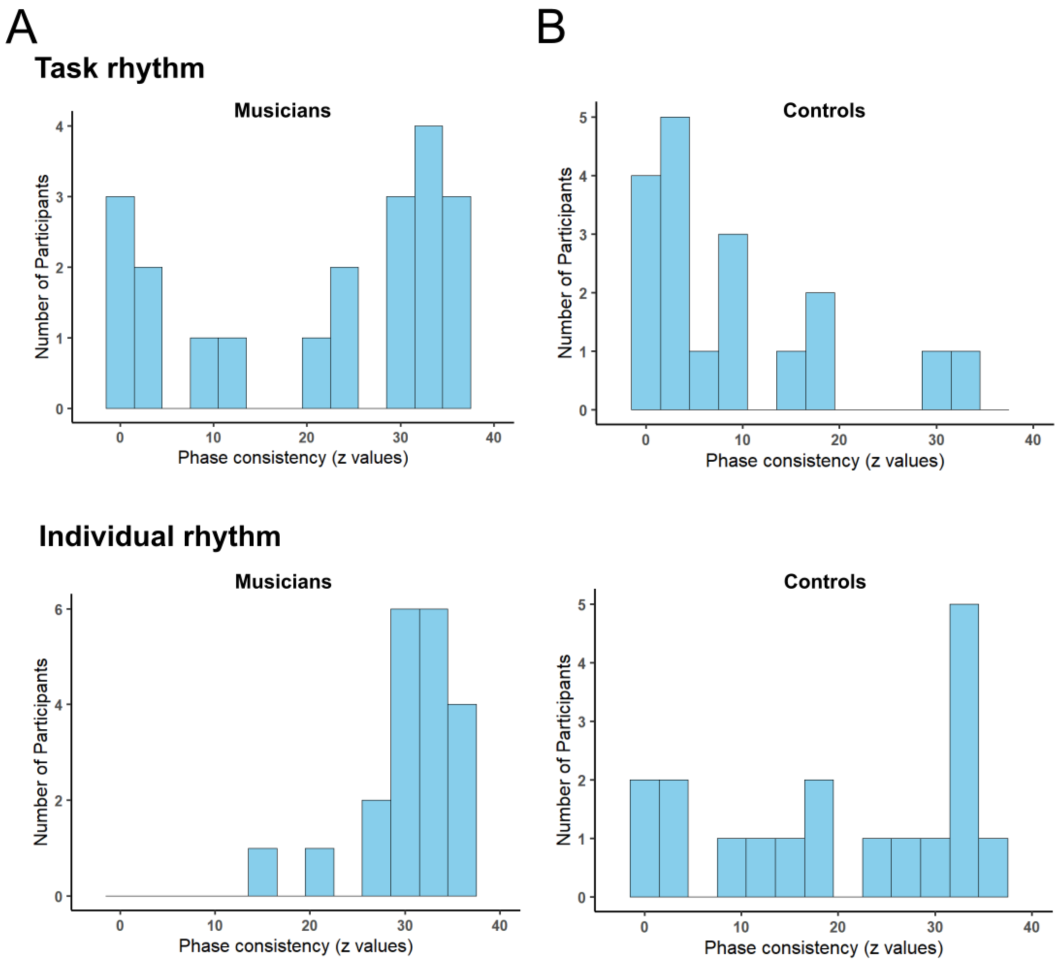


**S4 Fig. Distributions of phase consistency in the rhythmic task.** Histograms of *z*-statistic values from the Rayleigh test for musicians (A) and controls (B). For both analysis and visualization, the trial showing the best performance in reproducing the task rate (out of 2 trials) was selected for each participant.

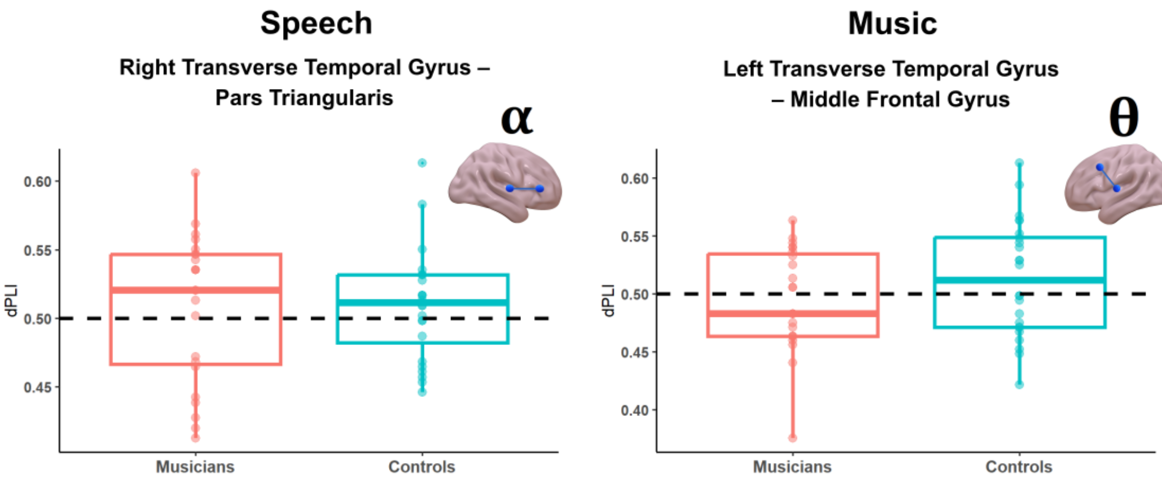


**S5 Fig. Results of directed connectivity analysis.** Boxplots of directed Phase Lag Index (dPLI) values. Horizontal dashed lines indicate the value reflecting no preference for either direction (dPLI = 0.5). Note that all values are between 0.4-0.6 and that about the same number of participants are below and above 0.5.

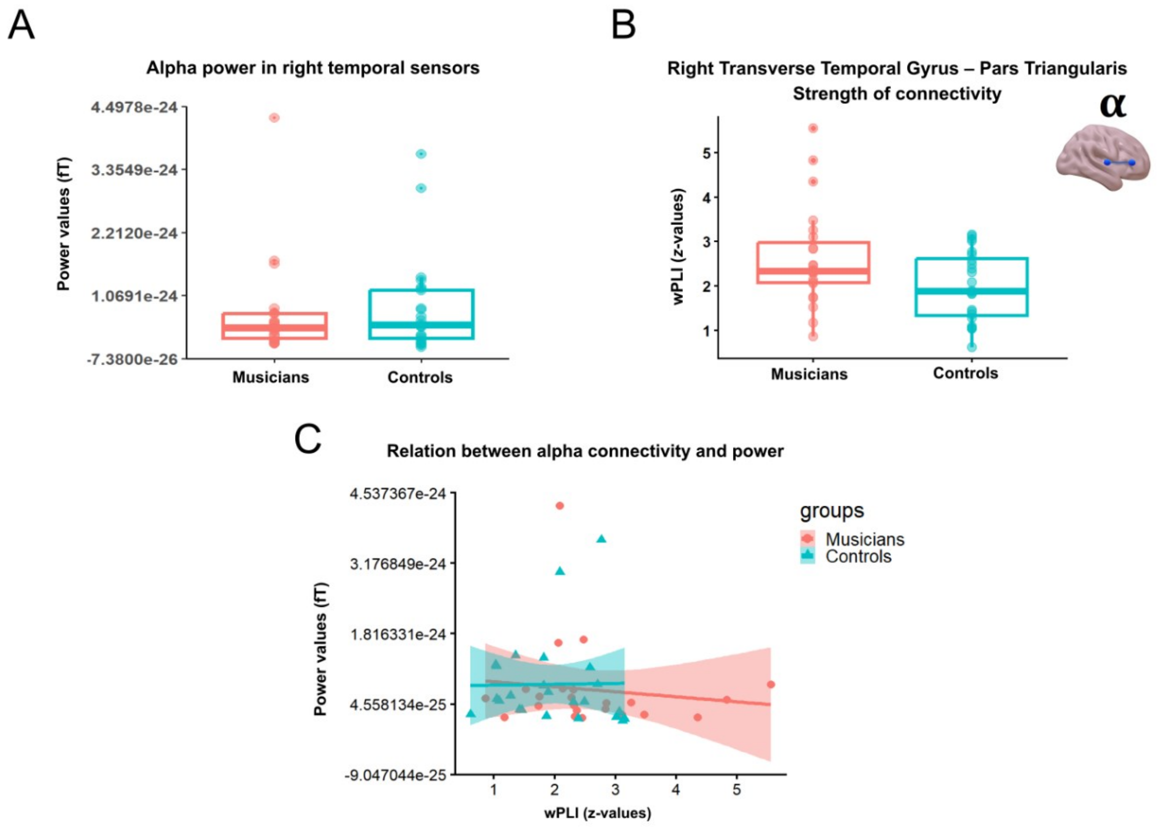


**S6 Fig. Alpha power and functional connectivity in the speech condition.** (A) Individual power values extracted at the same frequency bins used for functional connectivity. (B) Individual functional connectivity values (*z*-scored wPLI) used in linear regressions. (C) Relation between functional connectivity and power. Dots (musicians) and triangles (controls) represent individual participants.

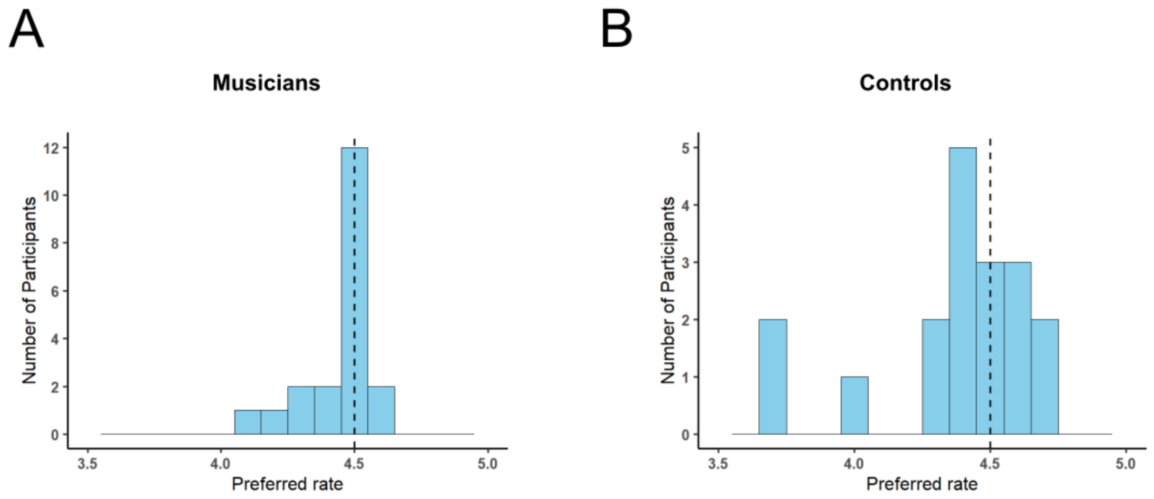


**S7 Fig. Distributions of preferred rates in the rhythmic task.** Vertical dashed lines indicate the task rate (4.5 Hz), which participants were instructed to repeat. Shown preferred rates were derived from the average Inter-Onset Interval (IOI) produced by participants in the trial showing the best performance in reproducing the task rate. Preferred rates and corresponding IOIs were used to compute phase consistency relative to the individual rhythm. (A) Musicians. (B) Controls.

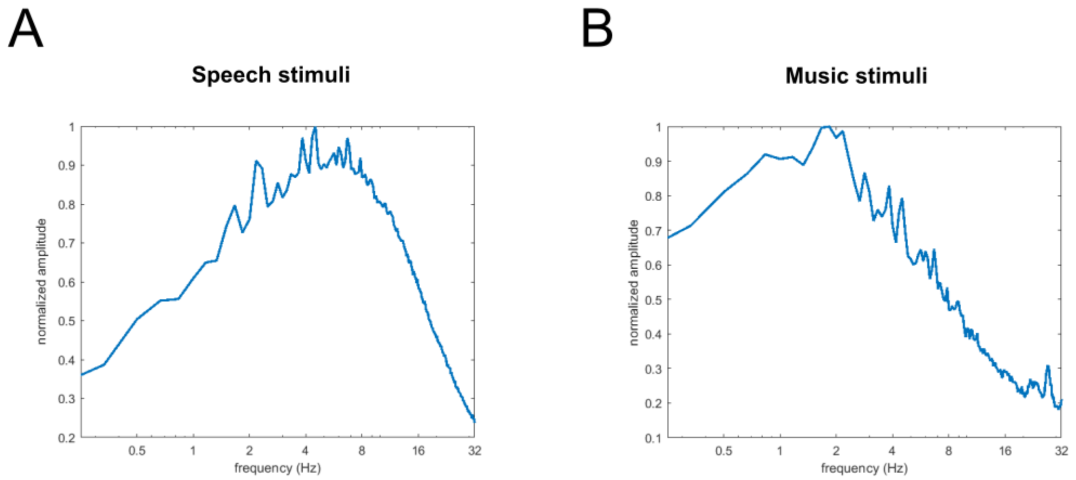


**S8 Fig. Modulation spectrum of stimuli.** Stimuli were concatenated and modulation spectrum was calculated on a logarithmic scale following the procedure described in Ding et al. [58]. (A) Speech stimuli. (B) Music stimuli.

**A**

| ID | Gender | Age | Number of instruments played | Primary instrument | Years of experience | Favourite genre |
|---|---|---|---|---|---|---|
| 1 | M | 52 | 1 | Piano | 44 | Classical music |
| 2 | F | 61 | 5+ | Piano | 55 | Classical music |
| 3 * | M | 55 | - | - | - | - |
| 4 | F | 51 | 4 | Accordion | 44 | Classical music |
| 5 | M | 49 | 1 | Accordion | 35 | Classical music |
| 6 | M | 53 | 1 | Trumpet | 42 | Classical music |
| 7 | M | 63 | 1 | Guitar | 44 | Jazz |
| 8 | M | 24 | 5+ | Trombone | 18 | Jazz |
| 9 | M | 22 | 2 | Transverse flute | 14 | Rock/pop |
| 10 | F | 22 | 5+ | Piano | 13 | Classical music |
| 11 | F | 22 | 3 | Piano | 14 | Classical music |
| 12 | F | 23 | 2 | Clarinet | 14 | Classical music |
| 13 | F | 23 | 3 | Transverse flute | 15 | Rock/pop |
| 14 | F | 24 | 3 | Clarinet | 15 | Classical music |
| 15 | F | 67 | 3 | Accordion | 10 | Rock/Pop |
| 16 | F | 60 | 4 | Piano | 50 | Jazz |
| 17 | M | 50 | 3 | Violin | 40 | Classical music |
| 18 | M | 36 | 2 | Violin | 30 | Rock/Pop |
| 19 | F | 48 | 2 | Cello | 40 | Classical music |
| 20 | M | 25 | 4 | Drums | 19 | Classical music |
| 21 | M | 55 | 2 | Txistu | 11 | Rock/Pop |
| 22 | F | 61 | 4 | Guitar | 31 | Rock/Pop |
| 23 | F | 50 | 3 | Piano | 40 | Classical music |
| 24 | F | 48 | 3 | Clarinet | 38 | Classical music |

**B**

| | ***PA*** | | ***AE*** | | ***SA*** | | ***EM*** | |
|---|---|---|---|---|---|---|---|---|
| | **M** | **SD** | **M** | **SD** | **M** | **SD** | **M** | **SD** |
| ***Musicians*** | 75.20 | 14.56 | 89.45 | 8.31 | 78.87 | 12.03 | 86.95 | 8.48 |
| ***Non-musicians*** | 51 | 15.75 | 69.22 | 14.93 | 50.22 | 17.51 | 74.13 | 16.97 |

**S1 Table. Characterization of participants' musical experience.** (A) Musical experience and demographics of musicians. (B) Percentage scores in subscales of the GOLD-MSI questionnaire. PA: Perceptive Abilities; AE: Active Engagement; SA: Singing Abilities; EM: Emotional engagement. A reduced version of the questionnaire was used, which included 31 items. The Musical Training subscale was not considered as training was covered in detail by the ELMEQ questionnaire.

| Title | Source |
|---|---|
| La bella y la bestia *(The beauty and the best)* | https://www.bosquedefantasias.com/recursos/cuentos-clasicos-populares/cuento-tradicional-la-bella-y-la-bestia |
| El dragón amigable *(The friendly dragon)* | https://www.bosquedefantasias.com/recursos/cuentos-dragones/cuento-el-dragon-amigable |
| El patito feo *(The ugly duckling)* | https://www.bosquedefantasias.com/recursos/cuentos-clasicos-populares/cuento-clasico-el-patito-feo |
| Paula la pirate *(Paula the pirate)* | https://www.bosquedefantasias.com/recursos/cuentos-piratas/paula-la-pirata |
| La bella durmiente *(The sleeping beauty)* | https://www.bosquedefantasias.com/recursos/cuentos-clasicos-populares/cuento-infantil-la-bella-durmiente |
| La magia del mar *(The magic of the sea)* | https://www.bosquedefantasias.com/recursos/cuentos-sirenas/la-magia-del-mar |
| Los tres cerditos *(The three piglets)* | https://www.bosquedefantasias.com/recursos/cuentos-clasicos-populares/cuento-clasico-infantil-los-tres-cerditos |
| El niño de la luna *(The child of the moon)* | https://www.bosquedefantasias.com/recursos/cuentos-para-dormir/el-nino-de-la-luna |
| Chopin – Nocturne Op. 9 No. 1 | https://www.youtube.com/watch?v=2bvg232HOn8 |
| Balakirev - Nocturne No.3 | https://www.youtube.com/watch?v=f2IY0_uKAF4&list=RDf2IY0_uKAF4&start_radio=1 |
| Tchaikovsky - The Seasons, Op. 37b: No. 6, June | http://youtube.com/watch?v=88y3sxpZ2ig |
| Rimsky-Korsakov - A Little Song | https://www.youtube.com/watch?v=dI3IlL0Trp0&list=RDdI3IlL0Trp0&start_radio=1 |
| Rachmaninoff - Rhapsody on a Theme by Paganini | https://www.youtube.com/watch?v=NsyTt587qeU |
| Dvořák - Theme and Variations, Op. 36: Theme | https://www.youtube.com/watch?v=a1pkCeFleF0&list=RDa1pkCeFleF0&start_radio=1 |
| Debussy - Arabesque No. 1 | https://www.youtube.com/watch?v=cVYH-7QGE-A |
| Fauré - Nocturne n. 3 in A flat major, op. 33 n. 3 | https://www.youtube.com/watch?v=DE_amYpTn9c&list=RDDE_amYpTn9c&start_radio=1 |

**S2 Table. Speech and music stimuli and online sources.**
